# Network remodeling induced by transcranial brain stimulation: A computational model of tDCS-triggered cell assembly formation

**DOI:** 10.1101/466136

**Authors:** Han Lu, Júlia V. Gallinaro, Stefan Rotter

**Affiliations:** Bernstein Center Freiburg & Faculty of Biology, University of Freiburg, Freiburg, Germany; Institute of Cellular and Integrative Neuroscience, University of Strasbourg, Strasbourg, France

## Abstract

Transcranial direct current stimulation (tDCS) is a variant of non-invasive neuromodulation, which promises treatment for brain diseases like major depressive disorder. In experiments, long-lasting aftereffects were observed, suggesting that persistent plastic changes are induced. The mechanism underlying the emergence of lasting aftereffects, however, remains elusive. Here we propose a model, which assumes that tDCS triggers a homeostatic response of the network involving growth and decay of synapses. The cortical tissue exposed to tDCS is conceived as a recurrent network of excitatory and inhibitory neurons, with synapses subject to homeostatically regulated structural plasticity. We systematically tested various aspects of stimulation, including electrode size and montage, as well as stimulation intensity and duration. Our results suggest that transcranial stimulation perturbs the homeostatic equilibrium and leads to a pronounced growth response of the network. The stimulated population eventually eliminates excitatory synapses with the unstimulated population, and new synapses among stimulated neurons are grown to form a cell assembly. Strong focal stimulation tends to enhance the connectivity within new cell assemblies, and repetitive stimulation with well-chosen duty cycles can increase the impact of stimulation even further. One long-term goal of our work is to help optimizing the use of tDCS in clinical applications.

## Introduction

Transcranial direct current stimulation (tDCS) is a non-invasive brain stimulation technique, where a weak constant current (1 − 2 mA) is applied to the brain via large electrodes attached to the scalp (Edwards et al., 2013). tDCS induces weak electric fields which are typically not sufficient to trigger action potentials directly, but can polarize the membrane of neurons by fractions of millivolts (Joucla and Yvert, 2009), depending on the orientation of the electric field vector relative to the somato-dendritic axis of the neuron (Wiethoff et al., 2014; Gluckman et al., 1996; Radman et al., 2009). This membrane potential deflection can influence spike timing and firing rates of neurons which are part of an active network (Bikson et al., 2006; Vöröslakos et al., 2018). Similar to other methods of neuromodulation, tDCS is claimed to have a potential for alleviating symptoms of certain brain diseases, such as major depressive disorder (Nitsche et al., 2009; Loo et al., 2012) or chronic pain (Garcia-Larrea, 2016; Ngernyam et al., 2015).

Although there is a record of promising applications of tDCS, both positive and negative outcomes have been reported in the literature (Horvath et al., 2015). Typical issues are due to insufficient sensitivity of measurements, or large inter-subject and intra-subject variability (Wiethoff et al., 2014). Positive evidence includes immediate changes of neural activity caused by tDCS, observed both in humans and in rodents. Positron emission tomography (PET) in humans revealed that tDCS can influence the activity of neurons in different brain regions (Lang et al., 2005), but the most affected region varies with electrode montage (Kuo et al., 2013), skull thickness (Opitz et al., 2015), individual geometry of cortex (Opitz et al., 2015), preexisting lesions (Minjoli et al., 2017), and other aspects. Systematic transcutaneous current stimulation experiments in rats (Vöröslakos et al., 2018) could establish quantitative relations between the externally applied current, the induced electric field, the associated membrane potential deflection, and the resulting firing rate change.

In addition to the instant impact on activity during stimulation, a sustained modulation of neural activity was also observed in humans after stimulation was turned off. Lasting aftereffects of tDCS, measured as motor evoked potentials (MEP) triggered by transcranial magnetic stimulation (TMS), were first reported by Nitsche and Paulus (2000), and later confirmed in motor cortex (Nitsche and Paulus, 2001) and somatosensory cortex (Matsunaga et al., 2004). Animal studies suggested that the elevated activity and excitability is not due to reverberating networks (Gartside, 1968a). Rather, changes in synaptic protein synthesis (Gartside, 1968b) point towards increased synaptic plasticity. In turn, blocking either brain-derived neurotrophic factor (BDNF) (Fritsch et al., 2010), NMDA receptors (Nitsche et al., 2003) or calcium channels (Monte-Silva et al., 2013) leads to a reduction of the stimulation-induced increments of the field potential in mice, or MEP in humans. Recent evidence suggests that multiple forms of plasticity are in fact contributing to tDCS aftereffects. Monte-Silva et al. (2013) observed that fast facilitation, or early-LTP (e-LTP), was induced after a single tDCS session (13 min) and lasted for at least 2 h post stimulation. In contrast, 26 min stimulation resulted in a reduced MEP amplitude. More interestingly, repetitive tDCS with 20 min pauses interspersed (13 − 20 − 13 min) resulted in late facilitation, or late-LTP (l-LTP). An elevated MEP was observed one day after the second stimulation, but not immediately after it. Functional LTP-like plastic changes of existing synapses were observed in DCS (Ranieri et al., 2012). Given the time scales of l-LTP, structural plasticity involving network remodeling also seems to play a role for the aftereffects. Structural changes at a slower time scale, however, can easily be underestimated due to difficulties measuring synapse turnover and changes in neuronal morphology *in vivo*. In summary, it is likely that both Hebbian and homeostatic, as well as functional and structural forms of plasticity underlie tDCS aftereffects.

Quantitative models of network remodeling have previously been described in the literature. Butz et al. (2014) first introduced the term *homeostatic structural plasticity* with reference to previously published versions of the theory (Butz et al., 2009; Butz and van Ooyen, 2013; Van Ooyen, 2011), which was based on ample experimental evidence that structural plasticity (Trachtenberg et al., 2002; Oray et al., 2004; Holtmaat and Svoboda, 2009; Pfeiffer et al., 2018) as well as homeostatic regulation of activity (Turrigiano and Nelson, 2004; Lee et al., 2013; Keck et al., 2013) are constantly taking place in many brain areas. This homeostatic structural plasticity model was able to provide explanations for cortical reorganization after stroke (Butz et al., 2009) and lesion (Butz-Ostendorf and van Ooyen, 2017), and for the formation of certain global network features during development (Butz et al., 2014; Gallinaro and Rotter, 2018). In this model, changing the number of synaptic contacts between two neurons leads to an apparent facilitation or depression of this specific connection, and the model may therefore also account for some cases of functional plasticity. Based on these previous insights it seemed natural to explore the contribution of homeostatic structural plasticity to the long-lasting aftereffects of transcranial brain stimulation.

In the present work, we hypothesize that employing proper stimulation protocols and adequate current strengths, tDCS is potent enough to polarize single neurons in a network (Vöröslakos et al., 2018). Based on this assumption, we assess the effect of such membrane potential deflections on neuronal firing rates. In a neural network model with homeostatic structural plasticity, we then systematically explore the influence of various stimulation parameters known from tDCS practice, such as electrode size and montage, stimulation strength and repetitive stimulation protocols. Our results suggest that tDCS can indeed induce substantial network remodeling and cell assembly formation, and focused strong and/or repetitive stimulation with well-chosen duty cycles can effectively boost the connectivity of the cell assemblies formed. The enhanced cell assembly might contribute to the empirical finding of profound plastic responses and enhanced therapeutic effects observed in current tDCS applications with a high-definition montage (Kuo et al., 2013) and repetitive stimulation (Monte-Silva et al., 2013). Our analysis also provides explanations for some of the negative results in tDCS practice.

## Methods

### Neuron model

All large-scale simulations of plastic neuronal networks of this study were performed with the NEST simulator (Linssen et al., 2018). Most were simulated with NEST 2.14, while NEST 2.16 with MPI-based parallel computation was used in the long repetitive protocol to achieve long simulation times. The linear, current-based leaky integrate-and-fire (LIF) neuron model was used throughout. The dynamic behavior of this point neuron model is described by the ordinary differential equation

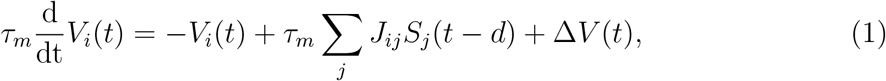

where *τ*_*m*_ is the membrane time constant. The variable *V*_*i*_(*t*) is the membrane potential of neuron *i*, with a resting value at 0 mV. ∆*V* (*t*) represents a polarization of the membrane imposed by an external electric field. The spike train generated by neuron *i* is denoted by 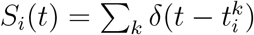, where 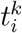 represent the individual spike times, and *d* is the synaptic transmission delay. The entries of the matrix *J*_*ij*_ denote the amplitude of the postsynaptic potential that is induced in neuron *i* upon the arrival of a spike from neuron *j*. In our model, all excitatory synapses have the amplitude *J*_*E*_ = 0.1 mV, whereas all inhibitory synapses have an amplitude of *J*_*I*_ = −0.8 mV. When the membrane potential *V*_*i*_(*t*) reaches the firing threshold, *V*_th_, an action potential is generated and the membrane potential is reset to *V*_reset_ = 10 mV. All parameters are once more listed in Table 1.

**Table 1:** Parameters of neuron model

| $\tau_m$ | $t_{\text{ref}}$ | $V_0$ | $V_{\text{reset}}$ | $V_{\text{th}}$ |
| --- | --- | --- | --- | --- |
| 10.0 ms | 2.0 ms | 0.0 mV | 10.0 mV | 20.0 mV |

### Model of transcranial DC stimulation

The electric field (EF) induced by tDCS can directly affect the membrane potential of neurons. Following Vöröslakos et al. (2018), we assumed that a strong enough EF will cause a small but significant membrane potential depolarization or hyperpolarization on some neurons in the network. The effective membrane potential deflection is determined by the orientation of the electric field vector relative to the somato-dendritic axis of the neuron (Wiethoff et al., 2014; Gluckman et al., 1996; Radman et al., 2009). When the electric field is properly aligned with the axis (apical dendrite closer to anode than soma), the somatic membrane potential is depolarized and the neuronal firing rate is increased. In contrast, if the electric field is perpendicular to the axis, it cannot influence the activity of this particular neuron. As a consequence, cells with extended and non-isotropic morphology, such as pyramidal neurons, should generally be more influenced by tDCS than the more compact inhibitory interneurons, which is also confirmed by Vöröslakos et al. (2018). Therefore, we assume only excitatory neurons to be sensitive to tDCS due to their spatial extent and non-isotropic morphology. We then asked the question, whether such a polarization could also cause significant changes in firing rate and, as a consequence, trigger structural plasticity and network remodeling. As our model neurons are actually point neurons with no spatial extent, we simply imposed an equivalent membrane potential bias ∆*V* on the soma of the neuron (Kayyali and Durand, 1991; Gluckman et al., 1996), see Figure 1A. This membrane potential bias also reflects the angle *θ* between the EF vector and somato-dendritic axis of the neuron with a factor cos(*θ*), see Figure 1B. The smallest magnitude of a membrane potential deflection reported in tDCS experiments to trigger physiological effects was in the range of 0.1 mV (Jackson et al., 2016; Vöröslakos et al., 2018).

**Figure 1:**
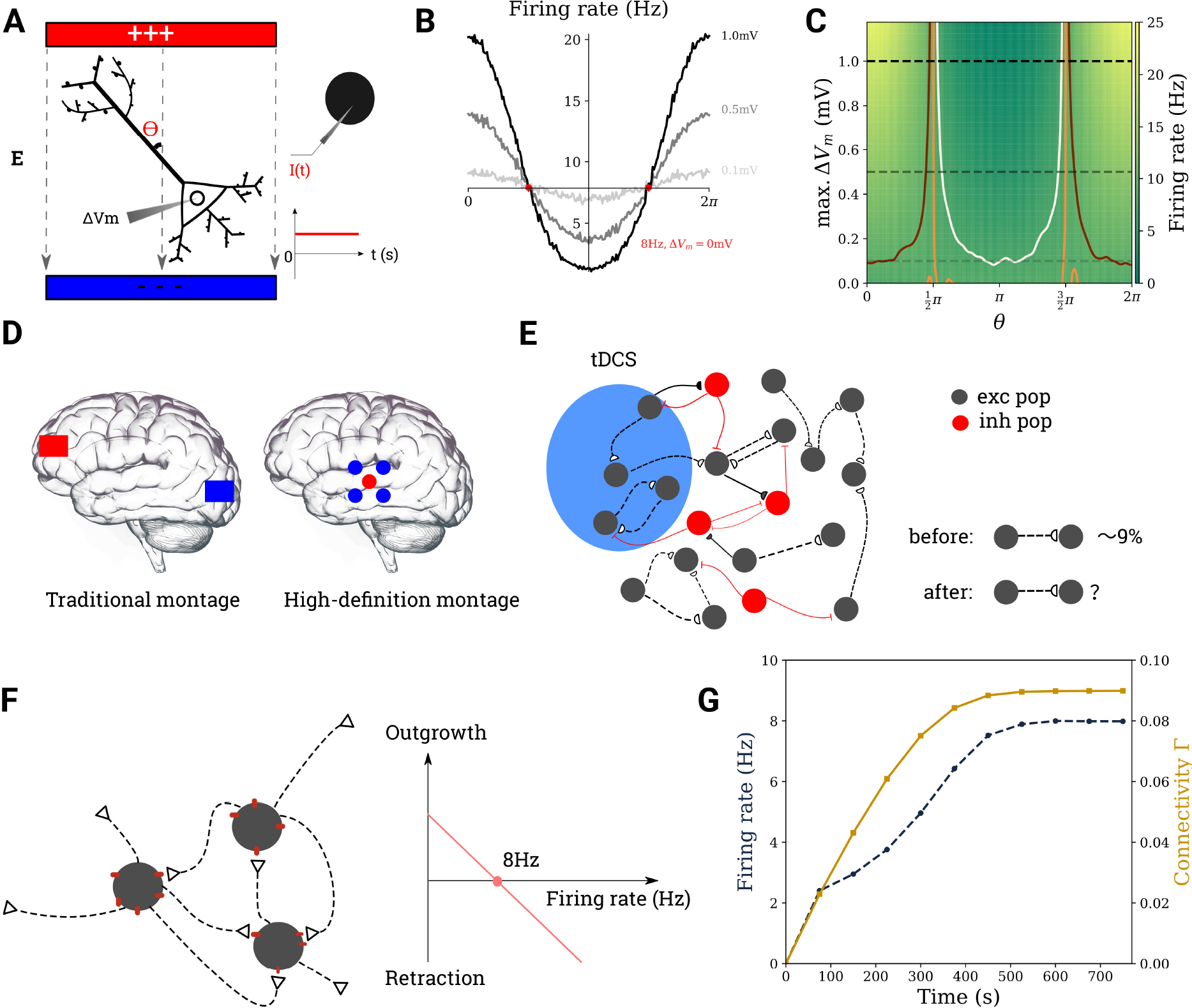
Modeling the effect of tDCS on cortical networks. **A** It is assumed that transcranial stimulation leads to a weak polarization of the neuron’s membrane potential (*left*). For a point neuron, this is achieved by injecting a current of suitable strength into its soma (*right*). **B** Firing rate modulation with the angle *θ* for 3 different values of ∆*V*_*m*_ (dotted lines on **C**). **C** Firing rate of a neuron the ongoing activity of which is modulated by tDCS, for different values of *θ* and membrane polarization ∆*V*_*m*_. The contour lines correspond to 7 Hz, 8 Hz, and 9 Hz in white, orange, and maron. **D** Electrode montages used in tDCS. **E** The region of interest subject to tDCS is modeled as a recurrent network of excitatory and inhibitory neurons. **F** Excitatory-to-excitatory synapses require the combination of a bouton (empty triangle) and a spine (red dot). The growth rate of both types of synaptic elements depends linearly on firing rate. **G** The network is grown from scratch before each tDCS stimulation experiment.

### Relative strength of background activity and tDCS

The effect of tDCS on a neuron with ongoing activity was assessed with single neuron simulations. The background input impinging onto the neuron was approximated by a spike train with Poisson statistics and rate *ν*_ext_ = 18.1 kHz, coupled to the neuron with synapses of strength *J*_ext_ = 0.1 mV. Given the parameters of our neuron model, this ongoing background activity leads to a fluctuating sub-threshold membrane potential with a mean value *µ* = *ν*_*ext*_*τ*_*m*_*J*_*ext*_ = 18.1 mV (Brunel, 2000). Different values of membrane polarization caused by tDCS (from 0.1 mV to 1.2 mV) were considered in our study, as described above. The firing rate of each condition was estimated from simulations of 100 s duration.

### Network model

Although there is a variety of EF distributions induced by different tDCS montages, we assume that neurons in the area most affected by stimulation are equally affected by tDCS (Jackson et al., 2016). This most affected area is modeled as an inhibition-dominated recurrent network (Brunel, 2000), comprising 10 000 excitatory and 2 500 inhibitory neurons. All connections involving inhibitory neurons were taken to be static. Excitatory and inhibitory synapses had fixed synaptic weights of *J*_*E*_ = 0.1 mV and *J*_*I*_ = − 0.8 mV, respectively. All these connections were randomly established, with 10% connection probability. In contrast, excitatory-to-excitatory (E-E) connections were subject to a growth rule called homeostatic structural plasticity (Gallinaro and Rotter, 2018; Butz and van Ooyen, 2013; Diaz-Pier et al., 2016). The network had initially no E-E connections whatsoever, and they were grown according to the specified rule during a growth period of 750 s for all simulations in the paper. Each neuron in the network received Poissonian external input at a rate of *r*_ext_ = 30 kHz. For the parameters chosen here, the network automatically entered an asynchronous-irregular state (Brunel, 2000). In all figures and simulations, transcranial DC stimulation was only applied after the end of the growth period. All network parameters are once more listed in Table 2.

**Table 2:**
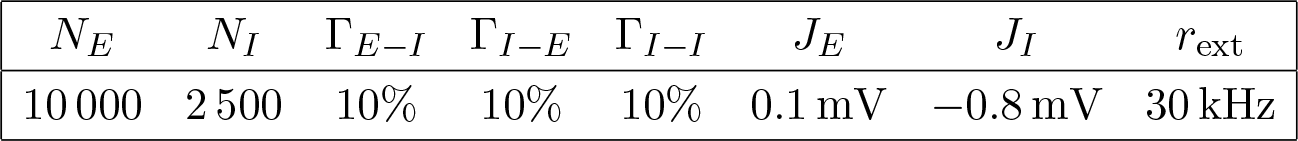
Parameters for network model

| $N_E$ | $N_I$ | $\Gamma_{E-I}$ | $\Gamma_{I-E}$ | $\Gamma_{I-I}$ | $J_E$ | $J_I$ | $r_{\text{ext}}$ |
| --- | --- | --- | --- | --- | --- | --- | --- |
| 10 000 | 2 500 | 10% | 10% | 10% | 0.1 mV | -0.8 mV | 30 kHz |

### Homeostatic structural plasticity

Connections between excitatory neurons underwent continuous remodeling, governed by rate-based homeostatic structural plasticity, as implemented in NEST (Diaz-Pier et al., 2016). Excitatory synapses were formed by combining a pre-synaptic element (bouton) and a post-synaptic element (spine). New synapses can form only if free synaptic elements are available. Pairs of neurons can form multiple synapses between them, and each individual functional synapse has the same weight *J*_*E*_ = 0.1 mV. It has been observed in experiments that neurite growth is governed by the concentration of intracellular calcium. It has been hypothesized that there is a set-point of the calcium concentration, which the neuron strives to reach and stabilize (Ramakers et al., 2001; Mattson and Kater, 1987). As a consequence, in the model of structural plasticity we use in our work, growth and deletion of synaptic elements are linked to the time-dependent intracellular calcium concentration *C*(*t*) = [Ca^2+^] of the neuron in question. In fact, this variable has been shown to be a good indicator of the neuron’s firing rate (Grewe et al., 2010). Whenever the neuron emits a spike, the intracellular calcium concentration experiences an increase by the amount *β*_Ca_ through calcium influx. Between spikes, the calcium concentration decays exponentially with time constant *τ*_Ca_,

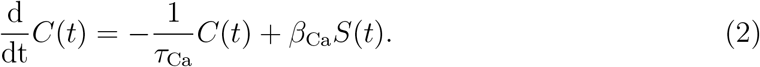

The synaptic growth rule is as follows. When the firing rate (or calcium concentration) falls below its set-point, the neuron will grow new synaptic elements and form functional synapses to compensate for the lack of excitatory input. In contrast, if the firing rate rises above the set-point, existing synapses are broken up and synaptic elements are removed. The respective counter-parts are added to the pool of free synaptic elements. We adopted a linear growth rule applying to both presynaptic and postsynaptic elements alike (Gallinaro and Rotter, 2018)

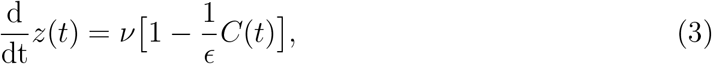

where *z*(*t*) is the total number of (presynaptic or postsynaptic) elements a neuron has available, *ν* is the growth rate, and *ϵ* is the target level of calcium concentration. In any given moment, free synaptic elements are randomly combined with matching free synaptic elements of other neurons, forming new functional synapses. All the parameters defining the structural plasticity rule are listed in Table 3.

**Table 3:** Parameters for the structural plasticity model

| $\epsilon$ | $\nu$ | $\tau_{\text{Ca}}$ | $\beta_{\text{Ca}}$ |
| --- | --- | --- | --- |
| 0.008 | $0.004 \text{ s}^{-1}$ | 10 s | 0.0001 |

### Protocols of transcranial DC stimulation

As suggested by current tDCS practice, many factors are essential to the outcome of a stimulation. For example, the traditional montage of two large sponge electrodes of size 5 cm × 7 cm induces a diffusive and weak EF. In contrast, high-definition montage using a small anodal electrode surrounded by several small cathodal electrodes induces a focal and relatively strong EF for the same stimulation current (Edwards et al., 2013). High-definition montage induces higher current densities, affects smaller populations, and possibly opposite field polarity at the edge of the cathodes. This method also exhibits better performance in tDCS practice, as compared to conventional montage (Kuo et al., 2013). To test these factors in our model we employed three different scenarios and systematically changed the size of the stimulated focus and the intensity of the stimulation in all of them. A summary of the parameters used in the different stimulation protocols described in this section can be found in Table 4.

**Table 4:** Configurations of DC stimulation

| Figure | Protocol | $f_{G1}$ | $\Delta V_1$ [mV] | $f_{G2}$ | $\Delta V_2$ [mV] | $f_{G3}$ | $\Delta V_3$ [mV] | Growth [s] | Repetition | $t_{stim}$ or $t1$ [s] | $t_{relax}$ or $t2$ [s] |
| --- | --- | --- | --- | --- | --- | --- | --- | --- | --- | --- | --- |
| 2B | uni-group | 10% | 0.1 | 90% | 0 | - | - | 750 | no | 150 | 300 |
| 2D | uni-group | 10% | 0.1 | 90% | 0 | - | - | 750 | no | 150 | 300 |
| 3A | tri-group | 30% | 0.1 | 30% | -0.1 | 40% | 0 | 750 | no | 150 | 300 |
| 3B | bi-group | 30% | 0.1 | 70% | -0.1 | - | - | 750 | no | 150 | 300 |
| 4A | bi-group | 10%, 30%, 50%, 70% | -1.2, 1.2<sup>1</sup> | 1 - $f_{G1}$ | -$\Delta V_1$ | - | - | 750 | no | 150 | 5850 |
| 4B | uni-group | 10%, 30%, 50%, 70% | -1.2, 1.2 | 1 - $f_{G1}$ | 0 | - | - | 750 | no | 150 | 5850 |
| 4C | bi-group | 10%, 20%, 30%, 40% | -1.2, 1.2 | $f_{G1}$ | -$\Delta V_1$ | 1 - $f_{G1}$ - $f_{G2}$ | 0 | 750 | no | 150 | 5850 |
| 4I | all | 50% | -1.2, 1.2 | 50% | -$\Delta V_1$ | - | - | 750 | no | 150 | 5850 |
| 5D | repetitive | 10% | 0.1 | 90% | 0 | - | - | 750 | yes<sup>2</sup> | multiple<sup>3</sup> |  |
| 5E | repetitive | 10% | multiple<sup>4</sup> | 90% | 0 | - | - | 750 | 80 | 75 | 150 |
| 6-on-off | repetitive | 10% | 0.1 | 90% | 0 | - | - | 750 | 3 | 150 | 150 |
| 6-alternating1 | repetitive | 10% | $\pm 0.05$ | 90% | 0 | - | - | 750 | 3 | 150 | 150 |
| 6-alternating2 | repetitive | 10% | $\pm 0.1$ | 90% | 0 | - | - | 750 | 3 | 150 | 150 |

<sup>1</sup> The stimulation intensities are -1.2, -0.8, -0.4, 0.4, 0.8, 1.2 mV
<sup>2</sup> The repetition round ( $n$ ) were matched with $n * t_1 = 6000s$
<sup>3</sup> The combinations used are (75, 75), (75, 150), (150, 75), (150, 150), (150, 300), (300, 150) s
<sup>4</sup> The stimulation intensities are 0.02, 0.04, 0.06, 0.08, 0.1, 0.2, 0.3, 0.4, 0.5 mV
All results in this study except Figure 4I and Figure 5E are averages from 30 independent simulations.

#### Uni-group

The first protocol we considered was a simplified scenario, in which only a subgroup of excitatory neurons in a large network was polarized by tDCS according to the above described protocol, while the remaining neurons were not affected and received only baseline external input. The focality of the stimulation is quantified by the percentage of excitatory neurons stimulated *f*_*G*1_. The more focused a stimulation is, the smaller is the subgroup of neurons affected by tDCS. The intensity of stimulation, on the other hand, is quantified by the amount of polarization. A stronger EF would lead to stronger membrane polarization of the soma of the model neurons. After a certain stimulation time *t*_stim_, tDCS is turned off and the network is allowed to relax for a period of *t*_relax_. Table 4 shows the values of *f*_*G*1_ and ∆*V*, as well as *t*_stim_ and *t*_relax_, used for the different figures.

#### Bi-group

Neurons in biological brains may not be uniformly polarized by stimulation. This is reflected in the bi-group scenario, in which a subgroup of neurons containing a fraction *f*_*G*1_ of all excitatory neurons is polarized by ∆*V*_1_ (similarly to the uni-group scenario), while the remaining excitatory neurons *f*_*G*2_ are stimulated with the same magnitude, but opposite polarity ∆*V*_2_. Similarly to the uni-group scenario, after a certain stimulation time *t*_stim_, tDCS is turned off and the network is simulated for a relaxation period *t*_relax_. The effect of stimulation on connectivity *I*_*G*_ was calculated as described below.

#### Tri-group

We designed yet another protocol, the tri-group scenario, to study the interaction of two actively stimulated subgroups with an unstimulated background. Two subgroups of excitatory neurons of the same size *f*_*G*1_ and *f*_*G*2_ are stimulated with the same magnitude, but different polarity ∆*V*_1_ and ∆*V*_2_. The remaining excitatory neurons in the network *f*_*G*3_ remain unstimulated. The resulting effect of stimulation on connectivity is measured as described below.

#### Repetitive patterns

To examine the effects of repetitive on-off stimulation, a certain fraction *f*_*G*1_ of the excitatory neurons was stimulated in multiple cycles with the uni-group protocol. Each cycle corresponds to a stimulation period of length *t*1 followed by a pause of length *t*2. The number of cycles *n*_*c*_ in each scenario was arranged to achieve a total DC stimulation time of *n*_*c*_*t*_1_ = 6 000 s.

Repetitive alternating stimulation is similar to the repetitive on-off protocol based on the uni-group scenario. The difference is that, instead of pausing, neurons are stimulated with opposite polarity and same magnitude. In table 4 we compiled a summary of all parameters for the stimulation protocols considered in our study.

### Measurements and calculations

#### Firing rate

The firing rate of a neuron was calculated from its spike count, in a 5 s activity recording, unless stated otherwise. The mean firing rate of a population was taken to be the arithmetic mean of firing rates across neurons in the group.

#### Synaptic connectivity

Let (*A*_*ij*_) be the *n × n* connectivity matrix of a network with *n* neurons. Its columns correspond to the axons, its rows correspond to the dendrites of the neurons involved. The specific entry *A*_*ij*_ of this matrix represents the total number of synapses from the presynaptic neuron *j* to the postsynaptic neuron *i*. The mean connectivity of this network is then given by 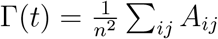, where *t* is the observing time point.

#### Time integral of the connectivity

When comparing the effects of different stimulation scenarios, one cannot simply consider the connectivity of the cell assembly at the end of simulation, because connectivity typically decays with certain time constants. We used the integrated connectivity change as a robust measure for the accumulated outcome of a stimulation. To account for the integrals, we first fit the connectivity change during the relaxation phase by a sum of three exponential decay functions

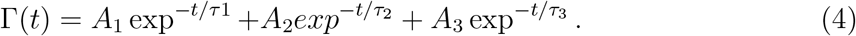

The parameter *A*_*k*_ is the amplitude of a component that decays with time constant *τ*_*k*_. We then computed the total integral of the connectivity by integrating the sum of exponentials, amounting to *I*_*G*_ = ∑_*k*_ *A*_*k*_*τ*_*k*_. This way we can also account for connectivity transients that persist for a very long time, extrapolating beyond the duration of our simulations.

## Results

### Immediate firing rate modulation by transcranial DC stimulation

We assume that the direct current applied to the brain during transcranial stimulation induces small deflections of the somatic membrane potential of neurons (Vöröslakos et al., 2018) and study the consequences of this deflection on neuronal firing rates. A polarization of the membrane ∆*V*_*m*_ in the range between −1.2 mV and 1.2 mV, which is not strong enough to elicit spikes in a neuron at rest, can nevertheless induce appreciable firing rate changes in a neuron with ongoing activity. Figure 1B and C shows how the firing rate of a model neuron driven by background input is modulated by both the strength of the depolarization and the angle *θ* between the electric field (EF) and the somato-dendritic axis. Even for a polarization as weak as ±0.1 mV, which is about the weakest depolarization known to cause observable physiological effects in tDCS experiments (Vöröslakos et al., 2018), the firing rate change was found to be larger than ±10% (Figure 1B, light gray curve). This suggests very clearly that tDCS can have an appreciable impact on neuronal activity, even if the stimulation intensity is apparently sub-threshold. As neuronal spiking can affect synaptic connectivity via activity-dependent plasticity, this raises the question whether transcranial stimulation can trigger plastic effects as well. To find an answer to this question, we set up a plastic network representing the tissue most affected by tDCS (Figure 1E-G) and study the effect of stimulation.

### Network remodeling triggered by transcranial DC stimulation

Different electrode montages are used in tDCS (Figure 1D), and they are thought to trigger different electric field distributions in the whole brain. We only modeled the most affected region stimulated by the peak current intensity. To explore the homeostatic response of the network, and the plastic processes associated with it, we first considered a simplified setting. In the uni-group scenario, only a subset of excitatory neurons in a larger network is stimulated (blue region in Figure 1E and Figure 2A). As shown in Figure 1F, tDCS disrupts the homeostatic equilibrium of the stimulated neurons by increasing their firing rate, initially leading to a deletion of synapses between stimulated neurons (see Methods for details of the structural plasticity model). When the stimulation has ceased, the firing rate of stimulated neurons drops due to a lack of recurrent input (Figure 2B), and the homeostatic process now triggers the formation of new synapses, predominantly among the stimulated neurons (Figure 2C). Figure 2F illustrates the process of cell assembly formation, similarly to what has been described previously by Gallinaro and Rotter (2018). Before and after the stimulation, assuming equilibrium in both cases, each neuron receives the same external input and fires at its target-rate (here, 8 Hz). Thus, the total number of input synapses from excitatory neurons will not have changed through stimulation. What has changed, however, is the source of input synapses: Before stimulation, input comes from both groups of neurons—to be stimulated (blue) and background (empty)—without any bias. During stimulation, however, synapses are broken up, and when stimulation is turned off, the stimulated neurons have more free synaptic elements to offer. Background neurons, which are only indirectly affected by stimulation and deviate less from their target rate, can only offer few synaptic elements to form new connections. Since the formation of new synapses is based on the availability of free elements, this leads to a bias for connections to be formed among stimulated neurons.

**Figure 2:**
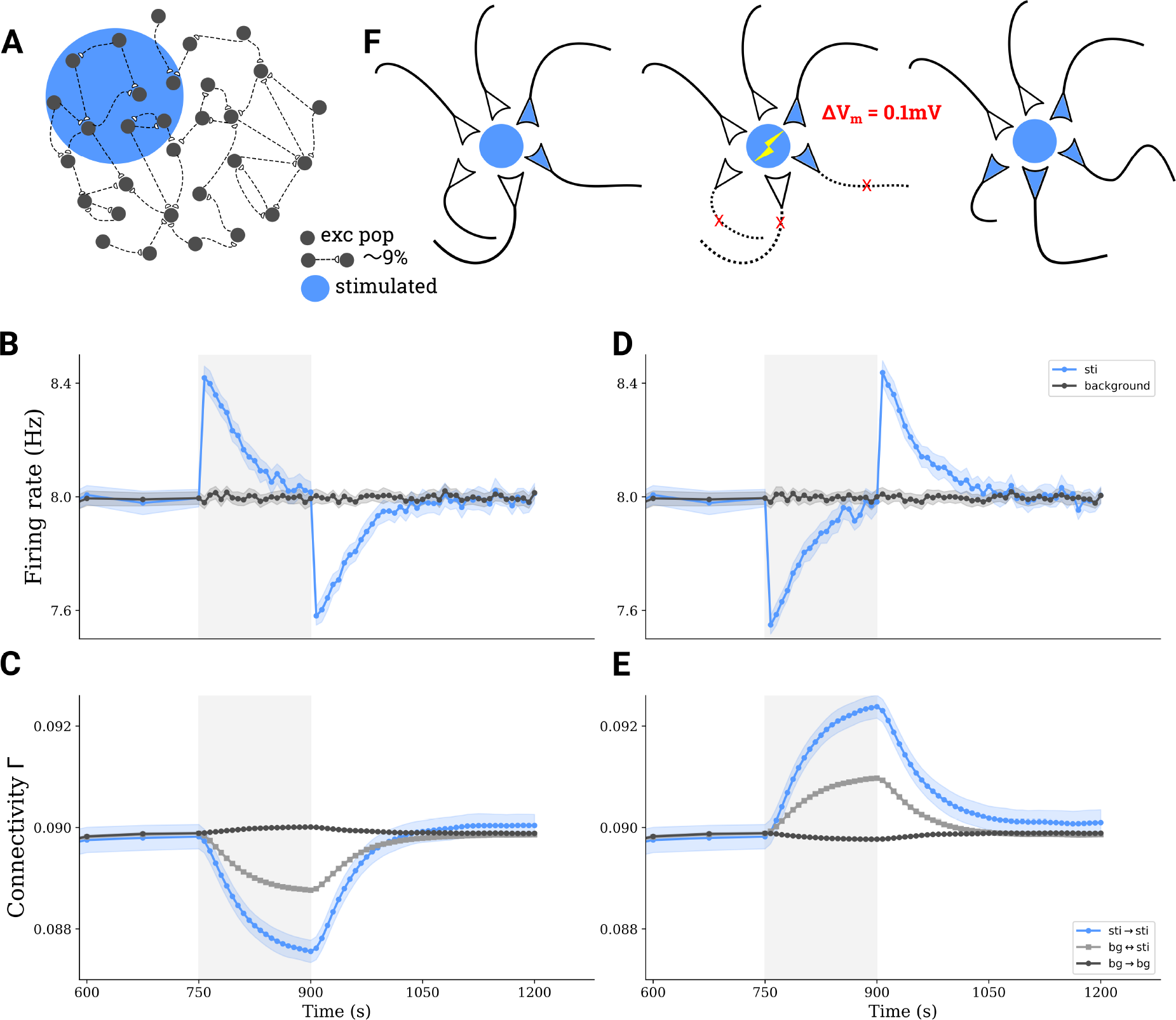
tDCS triggers the formation of cell assemblies. **A** A subgroup comprising 10% of all excitatory neurons in a larger network is stimulated by tDCS. **B** Average firing rate of directly stimulated (blue) and unstimulated (gray) excitatory neurons before, during and after applying a depolarizing stimulus. **C** Average connectivity among stimulated neurons (blue), among unstimulated neurons (dark gray), and between neurons belonging to different groups (light gray) upon depolarizing stimulation. **D-E** Similar to **B-C**, but for a hyperpolarizing stimulus. Shaded areas on **B-E** indicate the stimulation period. **F** Illustration explaining the process of structural plasticity that happened after a depolarizing tDCS. The stimulation triggers the removal of inter-population synapses, and accelerates the growth of synapses among stimulated neurons, leading to the formation of cell assemblies.

A similar process happens for hyperpolarizing DC (Figures 2D and E). In this case, however, the connectivity among stimulated neurons increases during tDCS due to hyperpolarization and a resulting drop in firing rate. In summary, any perturbation to the equilibrium of the network firing rate dynamics, no matter whether it is depolarizing or hyperpolarizing, will trigger an increased synaptic turnover and network remodeling by deleting between-group synapses and forming new synapses within the stimulated group to form a cell assembly.

### The effect of montage, focality, and intensity of transcranial DC stimulation

Stimulation is able to induce cell assembly formation in the uni-group scenario, as illustrated in Figure 2. However, neurons affected by tDCS might not be uniformly depolarized or hyperpolarized. Parameters like stimulation montage, focality or intensity certainly influence the degree to which each neuron in the stimulated population is affected, and to what extent its membrane potential is depolarized or hyperpolarized. Therefore, we investigated two alternative stimulation scenarios that capture some of the complexities of neuron polarization in real tissue: tri-group stimulation (Figure 3A) and bi-group stimulation (Figure 3B), the details of which are described in the Methods section. Similarly to the simplest scenario illustrated in Figure 2, the stimulated neurons again form a cell assembly (Figure 3E and F) also under more general conditions.

**Figure 3:**
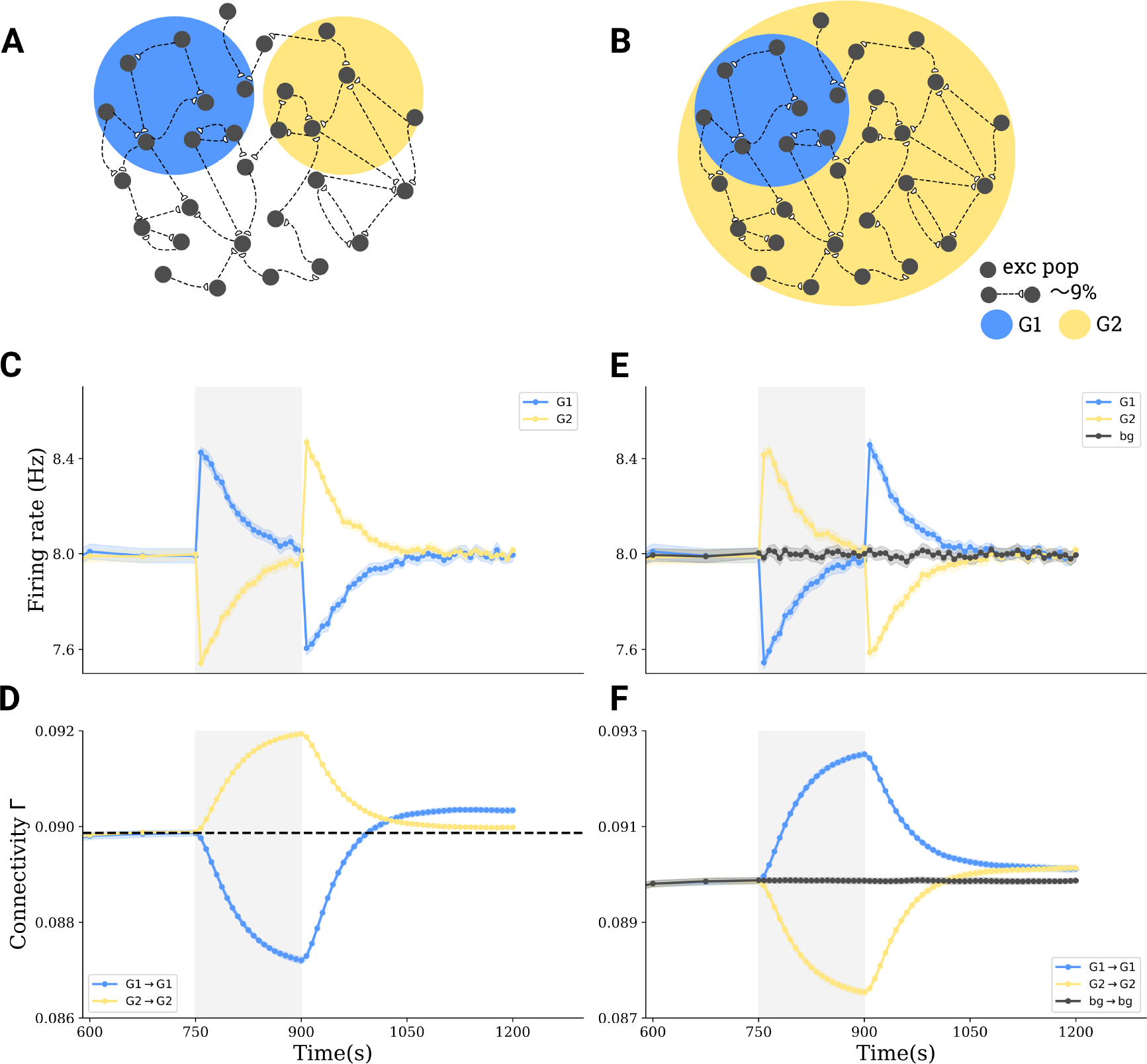
Interactions between sub-populations and cell assembly formation in more complex stimulation paradigms. **A** Tri-group scenario: 30% of all neurons in a network (G1) are depolarized by − 0.1 mV, another 30% (G2) are hyperpolarized by 0.1 mV, and the rest of 40% receives no stimulus. **B** Bi-group scenario: 30% (G1) are hyperpolarized by − 0.1 mV, and the remaining 70% (G2) are depolarized by 0.1 mV. **C,E** Group averages of firing rates in G1 (blue) and in G2 (yellow) before, during and after stimulation. **D,F** Group averages of the connectivity within G1 (blue), within G2 (yellow) and between G1 and G2 (gray).

We performed a systematic study covering different degrees of stimulation focality and intensity and compared the effects in all three scenarios: the bi-group (Figure 4A), uni-group (Figure 4B), and tri-group (Figure 4C). Higher stimulus intensity is implemented as a stronger membrane polarization, which results from a higher tDCS current density. Focality, quantified as the percentage of neurons in the network affected by membrane polarization, describes how focused stimulation is. More focused stimulation should have a polarizing effect on a smaller percentage of neurons. In each scenario, the connectivity in a newly formed cell assembly increases with absolute stimulation intensity and decreases with the size of the stimulated population (Figure 4D-F). We conclude that strong and focused stimulation (like high-definition stimulation) leads to stronger effects on the connectivity of the cell assembly. We further compared the effects of bi-group, uni-group and tri-group scenarios and found that the montage can greatly influence the outcome. When the polarization is very strong (above 0.8 mV) and focused, the effect *I*_*G*_1 is much stronger in the uni-group scenario as compared to the bi-group (Figure 4G) and tri-group (Figure 4H) scenario. But if the stimulus is weak, its effect in the bi-group scenario is larger than in the uni-group scenario. Therefore, using opposite polarities for stimulation could slightly boost cell assembly formation, provided the stimulus is weak. However, for strong and/or focused stimulation, uni-group stimulation leads to more pronounced cell assemblies.

**Figure 4:**
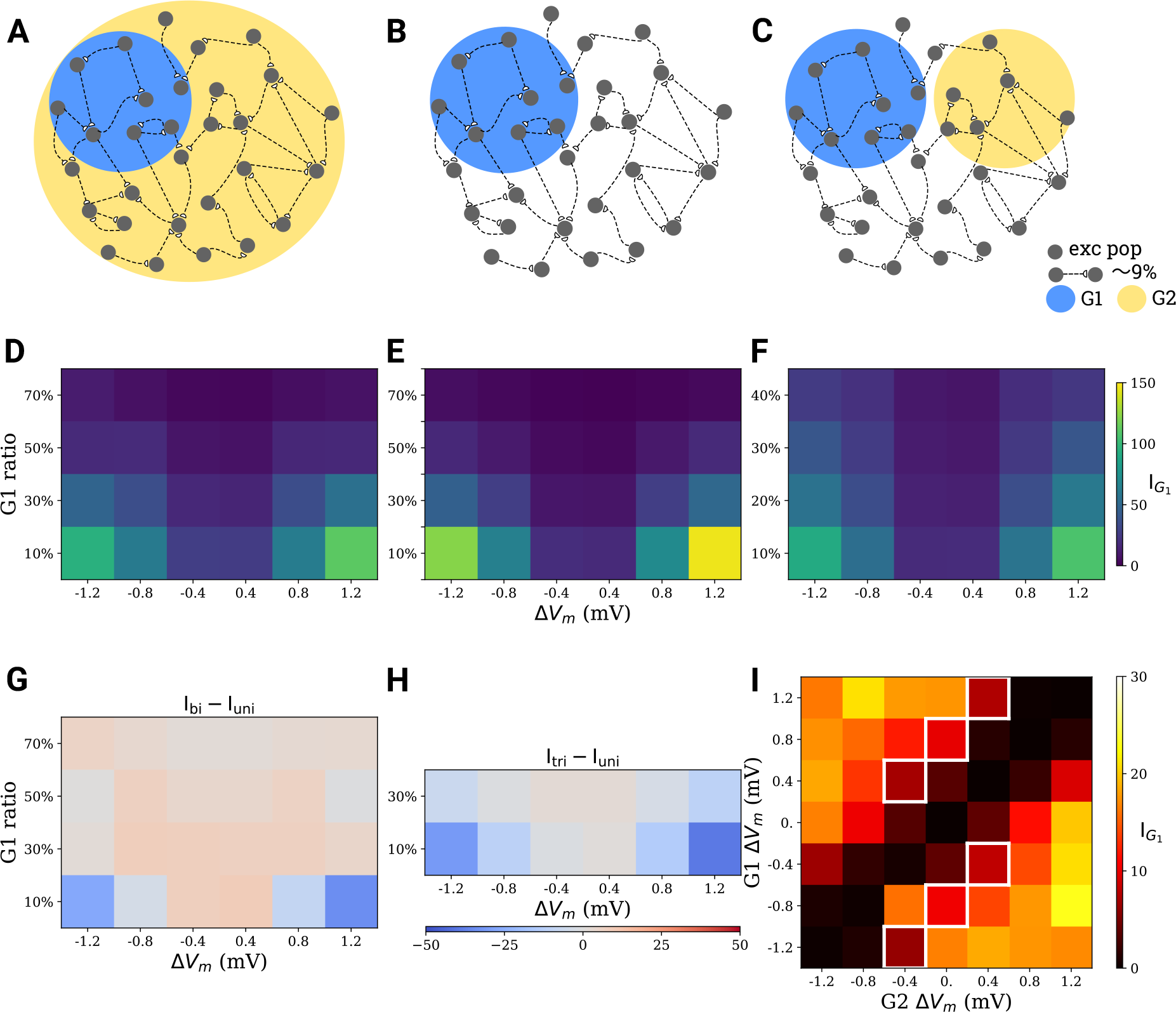
Comparison of tDCS effects with different electrode montage, as well as stimulus focality and intensity. **A** Bi-group stimulation scenario. **B** Uni-group stimulation scenario. **C** Tri-group stimulation scenario. **D-F** Integrated G1 cell assembly connectivity (*I*_*G*1_) at different focality and intensity levels for scenario **A-C**. **G,H** Difference between **D** and **E**, as well as **F** and **E**, respectively. **I** Integrated G1 cell assembly connectivity integrals (*I*_*G*1_) for different stimulation intensity levels for a specialized bi-group scenario, where G1 and G2 comprise half of the excitatory population, respectively. The white squares correspond to situations, where the difference between stimulation intensities of both groups amount to 0.8 mV.

The application of hyperpolarizing DC to all neurons in the background population can either amplify or attenuate the effect of the actual depolarizing stimulus. Two aspects might contribute to this phenomenon. Stimulating the background with reversed polarity increases the discrepancy of the stimulated group compared to the background (from ∆*V*_*m*_ to 2∆*V*_*m*_), but it may reduce the firing due to an activation of inhibitory neurons in the network. To disentangle the situation, we fixed the sizes of both the stimulated and the unstimulated group at 50% and then systematically changed the stimulus strength for both G1 and G2 in the range between −1.2 mV and 1.2 mV. The effect on G1 connectivity for different polarizations of G1 and G2 is displayed in Figure 4I. The values along the diagonal are very small, as there is no cell assembly formation when both groups experience the same stimulation. When the difference in stimulation of the two populations is large irrespective of its sign, the impact on G1 connectivity is also large (upper left and bottom right corners). We then checked, whether the relative difference between the polarization of G1 and G2 is sufficient to predict the stimulation outcome. The white squares in Figure 4I indicate simulations, in which the difference between G1 and G2 polarization is the same (0.8 mV), but the actual connectivities for individual groups are different. The strongest effect was achieved when the polarization of one of the two groups is 0 mV, which corresponds to the uni-group scenario. This supports the idea that network effects might influence the interaction between two groups, and that uni-group stimulation can achieve better outcomes than alterantive scenarios, provided stimulation is very strong.

### The effect of repetitive transcranial DC stimulation

Repetitive stimulation was simulated in our model by repeating stimulation of duration *t*_1_ in the uni-group scenario (Figure 2) multiple times, with a pause of duration *t*_2_ between successive stimulation periods (Figure 5A and B). The connectivity of the stimulated subpopulation generally increased upon repetition (Figure 5C). Figure 5D summarizes the outcome of different combinations of *t*_1_ and *t*_2_. Compared to long uninterrupted DC stimulus (single stimulation cycle with *t*_1_ = 6 000 s), repetitive stimulation (total stimulation time of 6 000 s distributed multiple cycles of shorter duration *t*_1_) led to higher final connectivity. We found that repetitive stimulation generally potentiated the effect of tDCS on cell assembly connectivity. Figure 5E demonstrates that after multiple repetitions, the connectivity appears to saturate at a level that essentially depends on the imposed polarization. As a consequence, a single stimulation with weak intensity for very long time does not necessarily lead to high connectivity, while repetitive stimulation at high intensity may lead to (much) higher connectivity. In our model we also tried very strong stimulation, repeated for several rounds. This lead to a very high assembly connectivity and eventually also to a very high firing rate of the excitatory population. High firing rates, in turn, induced a strong homeostatic response of the network and fast deletion of synapses, putting the network in an unfavorable and somewhat pathological state (data not shown).

**Figure 5:**
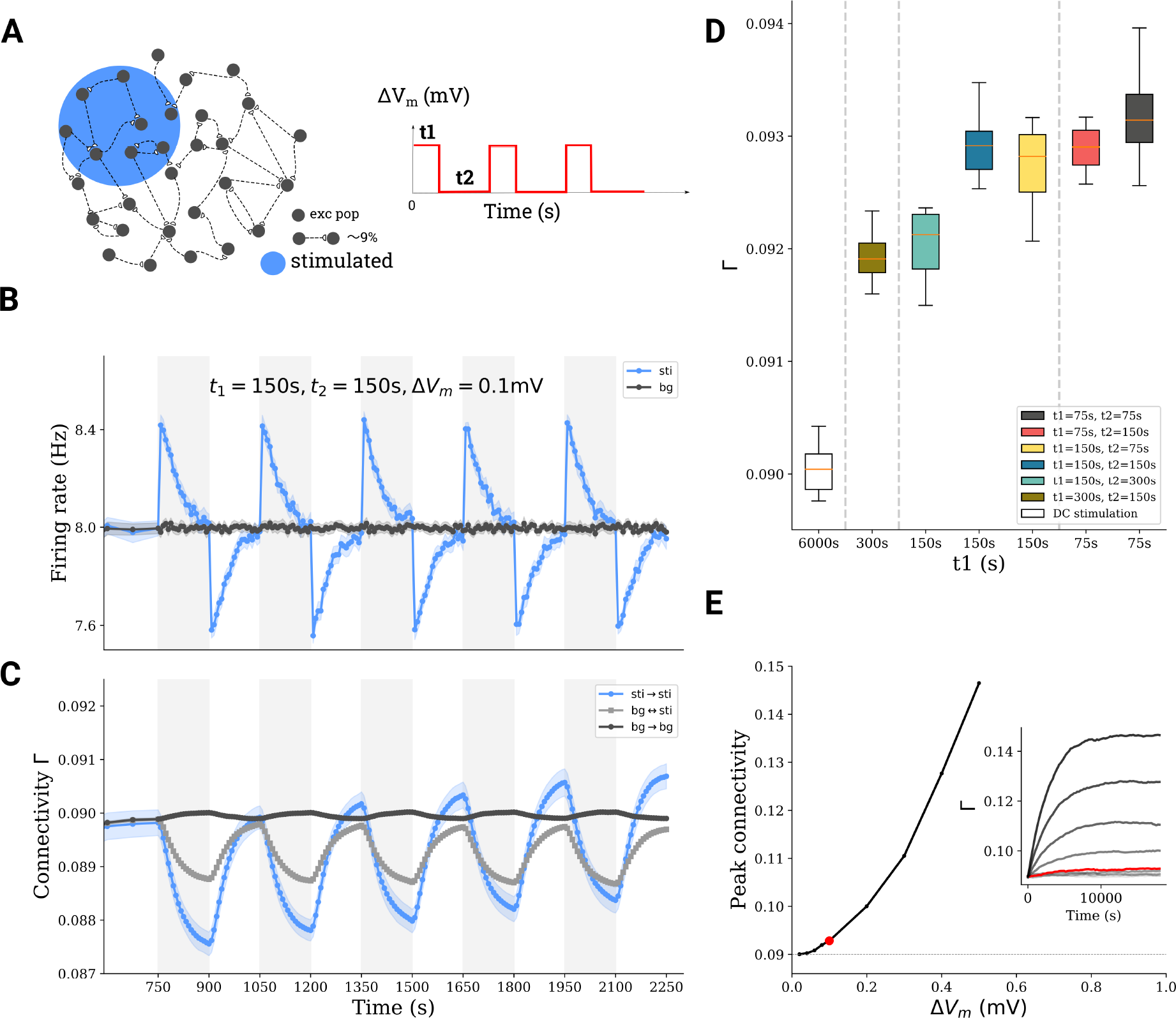
Repetitive stimulation boosts network remodeling. **A** A subnetwork of excitatory neurons (10%) is stimulated with a train of DC stimuli. Stimulation time is *t*_1_, followed by a pause of duration *t*_2_. **B,C** Average firing rate and connectivity during a train of stimuli. **D** For the same total stimulation time (6 000 s), the boosting depends on the exact repetition protocol. **E** The peak connectivity reached depends on the stimulation intensity, an asymmetric repetitive protocol (*t*_1_ = 75 s, *t*_2_ = 150 s) was used for all simulations here.

Repetitive stimulation can also be performed in cycles of alternating polarities, instead of a simple on-off protocol. Figure 6 shows the connectivity changes for two stimulation patterns: *on-off*, in which periods of depolarizing stimulation are followed by periods of no stimulation, and *alternating*, in which periods of depolarization are followed by periods of hyperpolarization. Simply substituting the off period by stimulation with different polarity seems to boost cell assembly connectivity (compare light green and dark brown traces in Figure 6B). However, if the alternating pattern has the same overall amplitude as the on-off stimulation (compare light brown and green traces in Figure 6B), the effect on cell assembly connectivity is the same as on the on-off pattern. Figure 6C depicts the final connectivity after 3 repetitions in 30 independent trials (mean and standard deviation are indicated in the inset).

**Figure 6:**
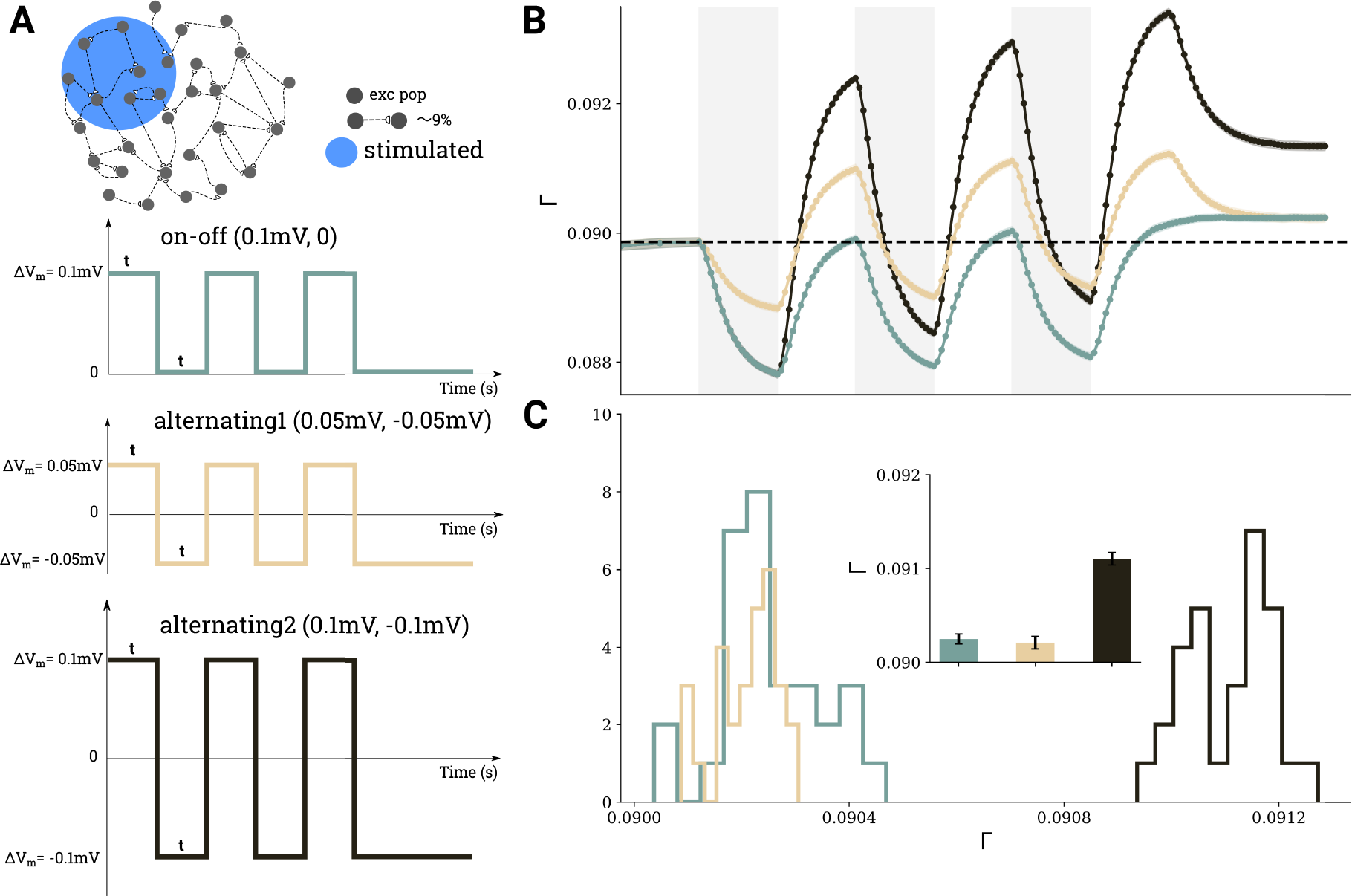
Comparison of three different scenarios for repetitive DC stimulation. **A** 10% of excitatory neurons were stimulated, using the same temporal protocol (*t*_1_ = 150 s, *t*_2_ = 150 s) in each case, but different amplitudes and polarities were employed, as indicated by the three different curves. **B** Evolution of average connectivity for the different stimulation scenarios, color code matches the stimulus curves in panel **A**. **C** Histograms of the connectivity reached after 3 cycles in the different scenarios extracted from 30 independent depolarizing simulations, mean values and standard deviations are shown in the inset.

## Discussion

We explored the plastic changes in network structure that can be induced by transcranial direct current stimulation (tDCS), exploiting the homeostatic response of synaptic growth and decay. We demonstrated that weak sub-threshold DC stimulation induces changes of neuronal firing rates and, thus, triggers network remodeling and cell assembly formation. Depolarized neurons first reduce the number of excitatory input synapses during stimulation, but then create new excitatory synapses predominantly with other stimulated neurons after stimulation is off. Interestingly, hyperpolarization also causes new synapses being formed preferentially among stimulated neurons. Stimulation triggers a profound and sustainable reorganization of network connectivity and leads to the formation of cell assemblies. With the help of our model, we explored different parameters of tDCS stimulation and found that strong and focused stimulation generally enhances the newly formed cell assemblies. We also observed that repetitive stimulation with well-chosen duty cycles can boost the induction of structural changes, and repetitive stimulation with alternating polarization may induce even higher connectivity changes.

We used network connectivity as a direct readout of stimulation effects, which is possible in model simulations, but cannot easily be done in experiments. However, the factors that we found to amplify the overall impact of stimulation are not unheard of in tDCS practice. Strong and focused stimulation, for example, which results from a high-definition electrode montage, does indeed lead to a stronger readout (MEP) and potentiates the therapeutic effects as compared to a conventional montage (Kuo et al., 2013). While applying the same total current, a high-definition montage induces stronger electric fields in smaller brain volumes (Edwards et al., 2013). Moreover, a high-definition montage narrows down the most affected brain region. We also found in our model that both factors indeed contribute to the induction of higher connectivity. Moreover, repetitive stimulation can boost connectivity, provided the duty cycles are chosen right. In fact, it has been demonstrated in experiments Monte-Silva et al. (2013) that two 13 min stimulations interrupted by a 20 min pause yields stronger MEP after-effects than a single, uninterrupted 26 min stimulation, while a repetition with a 24 hour pause in between could not accumulate the after-effects at all. In our model, we likewise found that multiple stimulation episodes with properly chosen pauses can achieve better effects than a single, uninterrupted stimulation.

Other computational approaches have been employed previously to analyze the neuron-scale mechanisms underlying tDCS or DCS. Most notably, Bikson et al. (2006) has explored several aspects of this: extracellular potassium concentration, polarization of the axonal terminal, action potential timing, and inhibitory neurons. Joucla and Yvert (2009) has provided an estimate of membrane potential changes for large axons exposed to an electric field, and Aspart et al. (2016) conceived the influence of the electric field on neuronal dendrites as external input to the soma. Another computational approach based on modern neural imaging methods has shed light on the question how strong the stimulation effects actually are. Spherical head models were first used to estimate the 3D current flow for any given electrode montage (Miranda et al., 2006). Later, fMRI based modeling was employed to devise individualized treatment of stroke or depressive patients (Datta et al., 2009; Ho et al., 2014; Huang et al., 2017). Our present work adopted insight and parameters from both approaches. In addition, we developed a new and original computational model to explore the impact of structural plasticity at the level of networks. This provides a bridge between the level of single neurons and the level of large-scale networks. Although our model contributes new explanations for some core observations in tDCS practice, there are still important issues left that cannot be appropriately addressed with our highly simplified model lacking relevant features of brain geometry. Also, the exact rules of growth and the time scales involved in homeostatic structural plasticity remain to be elucidated in experiments. To treat the influence of tDCS on network dynamics and structural plasticity of multiple brain regions would require a “network of networks” approach, which is, however, beyond the scope of our current study.

What are the actual effects of tDCS on network activity and function? Although robust and sustained effects of tDCS using relatively weak stimulation currents (1–2 mA) have been demonstrated (Nitsche et al., 2009; Nitsche and Paulus, 2000), Horvath et al. (2015) pointed to the difficulty reproducing positive results. Recently, Vöröslakos et al. (2018) have shown that the amount of membrane polarization due to tDCS depends on the strength of the applied current, and that there should be indeed no effect expected for very low intensities. Our simulation results suggest, however, that repetition could boost the impact on connectivity. The peak connectivity reached after sufficiently many repetitions, however, depends on stimulus intensity. Very weak stimulation cannot achieve high connectivity changes, even if repeated ad infinitum. Strong stimulation within a safe range could achieve higher connectivity, but too strong stimulation may lead to unfavorable network dynamics. Our model predicts very clearly that the accumulated effect achieved by stimulation depends not only on the exact repetition pattern, but also on stimulation intensity. On the other hand, a quantitative assessment of the aftereffects is difficult. In our work, the effect of tDCS on the network is quantified by measuring anatomical connectivity among stimulated and non-stimulated neurons. Such measurement is currently not possible in experiments, neither *in vivo* nor *in vitro*. Transcranial stimulation perturbs neuronal firing rates transiently and leads to the formation of cell assemblies, which persist after tDCS has been switched off and neuronal activity is back to baseline. Therefore, considering the homeostatic nature of structural plasticity, it is actually impossible to measure the effect of tDCS using simple neuronal activity measures. The question is, what are the effects of altered connectivity on the activity and the function of neuronal networks, and how can these effects be measured. This is a very interesting question, and the answer is complicated. Even if newly formed cell assemblies do not affect spontaneous activity as the firing rate of the neurons may be homeostatically regulated, they might still influence the evoked responses of neurons. Interestingly, Horvath et al. (2015) reviewed many tDCS studies and found that stimulation has a reliable effect only on the MEP amplitude, out of many potential biomarkers that were tested. The debate about the effects of tDCS on network function should, therefore, include the measures to quantify the outcome of a stimulation.

Another important issue raised by our work is that the total effect of stimulation might be too weak for detection. The connectivity changes triggered by a single cycle of polarization at ∆*V*_*m*_ = 0.1mV can only be detected if the full connectome is available for quantification. While possible in simulations, such a scenario is unrealistic in an experimental setting. Our simulation results suggest, however, that the outcome should increase upon repetitive stimulation and, therefore, possibly becomes easier to measure. The measurement time window of tDCS effects adds another puzzle to this question. The connectivity of the stimulated plastic network undergoes constant changes. During and after stimulation, for instance, total connectivity decreases and increases fast, constituting the homeostatic response. In contrast, the newly formed cell assembly persists for much longer periods and decays only with a slower time constant. It is not yet clear, however, which parameters influence this time constant, and it might be that different current intensity and electrode size have an impact on it. In fact, Jamil et al. (2017) recently observed in experiments that the current intensity might interact with the duration of stimulation needed for the homeostatic reversal of plasticity. If the exact stimulation protocol indeed influences the time scale of the aftereffect, naively comparing tDCS effects under different stimulation conditions “before” and “after” does not provide sufficient information regarding its outcome. In view of this, using a measure that takes the dynamics of the changes triggered by stimulation into account, such as the *I*_*G*_ measure introduced in this work, could quantify the effects of stimulation much more reliably.

In general, one needs to interpret the results and predictions of our work on network remodeling induced by tDCS with due caution. Our current work, however, could be a first step toward the goal of understanding and optimizing tDCS performance. More experiments addressing the impact of tDCS in human and in animal brains are definitely needed, and the results of our simulation study might indicate some new directions.

## Supportive Information

### Author contributions

The project was planned and realized by HL, JG, SR. JG established the network model with homeostatic structural plasticity. HL formalized the model of tDCS and performed all numerical simulations. HL analyzed the data; JR and SR contributed to the data analysis. HL wrote the manuscript, and all authors contributed to the revision.

## Code Accessibility

Simulation and analysis code is available upon request.

## Acknowledgments

This work is funded by the Universitätsklinikum Freiburg and NEUREX. Additional support by the German Research Foundation (DFG) through EXC 1086, and by the state of Baden-Württemberg through bwHPC and the German Research Foundation (DFG) through INST 39/963-1 FUGG is acknowledged. The authors thank Claus Normann, Lukas Frase, Andre Russowsky Brunoni, Sandra Diaz-Pier, and Benjamin Merkt for useful discussions. We also thank Uwe Grauer from the Bernstein Center Freiburg as well as Bernd Wiebelt and Michael Janczyk from the Freiburg University Computing Center for their assistance with HPC applications.

## References

Aspart, F., Ladenbauer, J., and Obermayer, K. (2016). Extending integrate-and-fire model neurons to account for the effects of weak electric fields and input filtering mediated by the dendrite. PLOS Computational Biology, 12(11):e1005206.

Bikson, M., Radman, T., and Datta, A. (2006). Rational modulation of neuronal processing with applied electric fields. In Engineering in Medicine and Biology Society, 2006. EMBS’06. 28th Annual International Conference of the IEEE, pages 1616–1619. IEEE.

Brunel, N. (2000). Dynamics of sparsely connected networks of excitatory and inhibitory spiking neurons. Journal of Computational Neuroscience, 8(3):183–208.

Butz, M., Steenbuck, I. D., and van Ooyen, A. (2014). Homeostatic structural plasticity increases the efficiency of small-world networks. Frontiers in Synaptic Neuroscience, 6:7.

Butz, M. and van Ooyen, A. (2013). A simple rule for dendritic spine and axonal bouton formation can account for cortical reorganization after focal retinal lesions. PLOS Computational Biology, 9(10):e1003259.

Butz, M., Van Ooyen, A., and Wörgötter, F. (2009). A model for cortical rewiring following deafferentation and focal stroke. Frontiers in Computational Neuroscience, 3:10.

Butz-Ostendorf, M. and van Ooyen, A. (2017). Is lesion-induced synaptic rewiring driven by activity homeostasis? In The Rewiring Brain, pages 71–92. Elsevier.

Datta, A., Bansal, V., Diaz, J., Patel, J., Reato, D., and Bikson, M. (2009). Gyri-precise head model of transcranial direct current stimulation: improved spatial focality using a ring electrode versus conventional rectangular pad. Brain Stimulation: Basic, Translational, and Clinical Research in Neuromodulation, 2(4):201–207.

Diaz-Pier, S., Naveau, M., Ostendorf, M., and Morrison, A. (2016). Automatic generation of connectivity for large-scale neuronal network models through structural plasticity. Frontiers in Neuroanatomy, 10:57.

Edwards, D., Cortes, M., Datta, A., Minhas, P., Wassermann, E. M., and Bikson, M. (2013). Physiological and modeling evidence for focal transcranial electrical brain stimulation in humans: a basis for high-definition tDCS. Neuroimage, 74:266–275.

Fritsch, B., Reis, J., Martinowich, K., Schambra, H. M., Ji, Y., Cohen, L. G., and Lu, B. (2010). Direct current stimulation promotes BDNF-dependent synaptic plasticity: potential implications for motor learning. Neuron, 66(2):198–204.

Gallinaro, J. V. and Rotter, S. (2018). Associative properties of structural plasticity based on firing rate homeostasis in recurrent neuronal networks. Scientific Reports, 8(1):3754.

Garcia-Larrea, L. (2016). tDCS as a procedure for chronic pain relief. Neurophysiologie Clinique/Clinical Neurophysiology, 46(3):224.

Gartside, I. B. (1968a). Mechanisms of sustained increases of firing rate of neurones in the rat cerebral cortex after polarization: reverberating circuits or modification of synaptic conductance? Nature, 220(5165):382.

Gartside, I. B. (1968b). Mechanisms of sustained increases of firing rate of neurones in the rat cerebral cortex after polarization: role of protein synthesis. Nature, 220(5165):383.

Gluckman, B. J., Neel, E. J., Netoff, T. I., Ditto, W. L., Spano, M. L., and Schiff, S. J. (1996). Electric field suppression of epileptiform activity in hippocampal slices. Journal of Neurophysiology, 76(6):4202–4205.

Grewe, B. F., Langer, D., Kasper, H., Kampa, B. M., and Helmchen, F. (2010). High-speed in vivo calcium imaging reveals neuronal network activity with near-millisecond precision. Nature Methods, 7(5):399.

Ho, K.-A., Bai, S., Martin, D., Alonzo, A., Dokos, S., Puras, P., and Loo, C. K. (2014). A pilot study of alternative transcranial direct current stimulation electrode montages for the treatment of major depression. Journal of Affective Disorders, 167:251–258.

Holtmaat, A. and Svoboda, K. (2009). Experience-dependent structural synaptic plasticity in the mammalian brain. Nature Reviews Neuroscience, 10(9):647.

Horvath, J. C., Forte, J. D., and Carter, O. (2015). Evidence that transcranial direct current stimulation (tDCS) generates little-to-no reliable neurophysiologic effect beyond MEP amplitude modulation in healthy human subjects: a systematic review. Neuropsychologia, 66:213–236.

Huang, Y., Liu, A. A., Lafon, B., Friedman, D., Dayan, M., Wang, X., Bikson, M., Doyle, W. K., Devinsky, O., and Parra, L. C. (2017). Measurements and models of electric fields in the in vivo human brain during transcranial electric stimulation. eLife, 6:e18834.

Jackson, M. P., Rahman, A., Lafon, B., Kronberg, G., Ling, D., Parra, L. C., and Bikson, M. (2016). Animal models of transcranial direct current stimulation: methods and mechanisms. Clinical Neurophysiology, 127(11):3425–3454.

Jamil, A., Batsikadze, G., Kuo, H.-I., Labruna, L., Hasan, A., Paulus, W., and Nitsche, M. A. (2017). Systematic evaluation of the impact of stimulation intensity on neuroplastic after-effects induced by transcranial direct current stimulation. The Journal of Physiology, 595(4):1273–1288.

Joucla, S. and Yvert, B. (2009). The “mirror” estimate: an intuitive predictor of membrane polarization during extracellular stimulation. Biophysical Journal, 96(9):3495–3508.

Kayyali, H. and Durand, D. (1991). Effects of applied currents on epileptiform bursts in vitro. Experimental Neurology, 113(2):249–254.

Keck, T., Keller, G. B., Jacobsen, R. I., Eysel, U. T., Bonhoeffer, T., and Hübener, M. (2013). Synaptic scaling and homeostatic plasticity in the mouse visual cortex in vivo. Neuron, 80(2):327–334.

Kuo, H.-I., Bikson, M., Datta, A., Minhas, P., Paulus, W., Kuo, M.-F., and Nitsche, M. A. (2013). Comparing cortical plasticity induced by conventional and high-definition 4 × 1 ring tdcs: a neurophysiological study. Brain Stimulation: Basic, Translational, and Clinical Research in Neuromodulation, 6(4):644–648.

Lang, N., Siebner, H. R., Ward, N. S., Lee, L., Nitsche, M. A., Paulus, W., Rothwell, J. C., Lemon, R. N., and Frackowiak, R. S. (2005). How does transcranial dc stimulation of the primary motor cortex alter regional neuronal activity in the human brain? European Journal of Neuroscience, 22(2):495–504.

Lee, K. J., Queenan, B. N., Rozeboom, A. M., Bellmore, R., Lim, S. T., Vicini, S., and Pak, D. T. (2013). Mossy fiber-CA3 synapses mediate homeostatic plasticity in mature hippocampal neurons. Neuron, 77(1):99–114.

Linssen, C., Lepperød, M. E., Mitchell, J., Pronold, J., Eppler, J. M., Keup, C., Peyser, A., Kunkel, S., Weidel, P., Nodem, Y., Terhorst, D., Deepu, R., Deger, M., Hahne, J., Sinha, A., Antonietti, A., Schmidt, M., Paz, L., Garrido, J., Ippen, T., Riquelme, L., Serenko, A., Kühn, T., Kitayama, I., Mørk, H., Spreizer, S., Jordan, J., Krishnan, J., Senden, M., Hagen, E., Shusharin, A., Vennemo, S. B., Rodarie, D., Morrison, A., Graber, S., Schuecker, J., Diaz, S., Zajzon, B., and Plesser, H. E. (2018). NEST 2.16.0.

Loo, C. K., Alonzo, A., Martin, D., Mitchell, P. B., Galvez, V., and Sachdev, P. (2012). Transcranial direct current stimulation for depression: 3-week, randomised, sham-controlled trial. The British Journal of Psychiatry, 200(1):52–59.

Matsunaga, K., Nitsche, M. A., Tsuji, S., and Rothwell, J. C. (2004). Effect of transcranial DC sensorimotor cortex stimulation on somatosensory evoked potentials in humans. Clinical Neurophysiology, 115(2):456–460.

Mattson, M. P. and Kater, S. B. (1987). Calcium regulation of neurite elongation and growth cone motility. Journal of Neuroscience, 7(12):4034–4043.

Minjoli, S., Saturnino, G. B., Blicher, J. U., Stagg, C. J., Siebner, H. R., Antunes, A., and Thielscher, A. (2017). The impact of large structural brain changes in chronic stroke patients on the electric field caused by transcranial brain stimulation. NeuroImage: Clinical, 15:106–117.

Miranda, P. C., Lomarev, M., and Hallett, M. (2006). Modeling the current distribution during transcranial direct current stimulation. Clinical Neurophysiology, 117(7):1623–1629.

Monte-Silva, K., Kuo, M.-F., Hessenthaler, S., Fresnoza, S., Liebetanz, D., Paulus, W., and Nitsche, M. A. (2013). Induction of late LTP-like plasticity in the human motor cortex by repeated non-invasive brain stimulation. Brain Stimulation: Basic, Translational, and Clinical Research in Neuromodulation, 6(3):424–432.

Ngernyam, N., Jensen, M. P., Arayawichanon, P., Auvichayapat, N., Tiamkao, S., Janjarasjitt, S., Punjaruk, W., Amatachaya, A., Aree-uea, B., and Auvichayapat, P. (2015). The effects of transcranial direct current stimulation in patients with neuropathic pain from spinal cord injury. Clinical Neurophysiology, 126(2):382–390.

Nitsche, M. A., Boggio, P. S., Fregni, F., and Pascual-Leone, A. (2009). Treatment of depression with transcranial direct current stimulation (tDCS): a review. Experimental Neurology, 219(1):14–19.

Nitsche, M. A., Fricke, K., Henschke, U., Schlitterlau, A., Liebetanz, D., Lang, N., Henning, S., Tergau, F., and Paulus, W. (2003). Pharmacological modulation of cortical excitability shifts induced by transcranial direct current stimulation in humans. The Journal of Physiology, 553(Pt 1):293–301.

Nitsche, M. A. and Paulus, W. (2000). Excitability changes induced in the human motor cortex by weak transcranial direct current stimulation. The Journal of Physiology, 527(3):633–639.

Nitsche, M. A. and Paulus, W. (2001). Sustained excitability elevations induced by transcranial dc motor cortex stimulation in humans. Neurology, 57(10):1899–1901.

Opitz, A., Paulus, W., Will, S., Antunes, A., and Thielscher, A. (2015). Determinants of the electric field during transcranial direct current stimulation. Neuroimage, 109:140–150.

Oray, S., Majewska, A., and Sur, M. (2004). Dendritic spine dynamics are regulated by monocular deprivation and extracellular matrix degradation. Neuron, 44(6):1021–1030.

Pfeiffer, T., Poll, S., Bancelin, S., Angibaud, J., Inavalli, V. K., Keppler, K., Mittag, M., Fuhrmann, M., and Nägerl, U. V. (2018). Chronic 2p-STED imaging reveals high turnover of dendritic spines in the hippocampus in vivo. eLife, 7:e34700.

Radman, T., Ramos, R. L., Brumberg, J. C., and Bikson, M. (2009). Role of cortical cell type and morphology in subthreshold and suprathreshold uniform electric field stimulation in vitro. Brain Stimulation: Basic, Translational, and Clinical Research in Neuromodulation, 2(4):215–228.

Ramakers, G., Avci, B., Van Hulten, P., Van Ooyen, A., Van Pelt, J., Pool, C., and Lequin, M. (2001). The role of calcium signaling in early axonal and dendritic morphogenesis of rat cerebral cortex neurons under non-stimulated growth conditions. Developmental Brain Research, 126(2):163–172.

Ranieri, F., Podda, M. V., Riccardi, E., Frisullo, G., Dileone, M., Profice, P., Pilato, F., Di Lazzaro, V., and Grassi, C. (2012). Modulation of LTP at rat hippocampal ca3-ca1 synapses by direct current stimulation. Journal of Neurophysiology, 107(7):1868–1880.

Trachtenberg, J. T., Chen, B. E., Knott, G. W., Feng, G., Sanes, J. R., Welker, E., and Svoboda, K. (2002). Long-term in vivo imaging of experience-dependent synaptic plasticity in adult cortex. Nature, 420(6917):788.

Turrigiano, G. G. and Nelson, S. B. (2004). Homeostatic plasticity in the developing nervous system. Nature Reviews Neuroscience, 5(2):97.

Van Ooyen, A. (2011). Using theoretical models to analyse neural development. Nature Reviews Neuroscience, 12(6):311.

Vöröslakos, M., Takeuchi, Y., Brinyiczki, K., Zombori, T., Oliva, A., Fernéndez-Ruiz, A., Kozak, G., Kincses, Z. T., Ivényi, B., Buzséki, G., et al. (2018). Direct effects of transcranial electric stimulation on brain circuits in rats and humans. Nature Communications, 9(1):483.

Wiethoff, S., Hamada, M., and Rothwell, J. C. (2014). Variability in response to transcranial direct current stimulation of the motor cortex. Brain Stimulation, 7(3):468–475.

