## supplementary-figures for "Network remodeling induced by transcranial brain stimulation: A computational model of tDCS-triggered cell assembly formation"

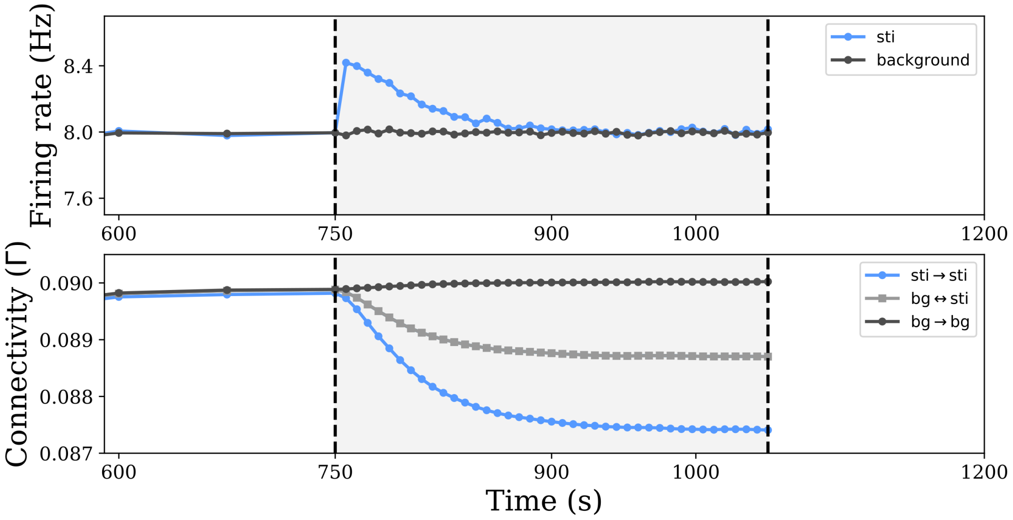


Figure S1: Changes in firing rate and connectivity over time, when a subgroup (10% of all excitatory neurons) is experiencing 2.5 pA DC stimulation. When the firing rate of the stimulated neurons reaches the set-point, the connectivity does not change any more, even if the stimulation is continued. The networks has then reached its structural equilibrium.

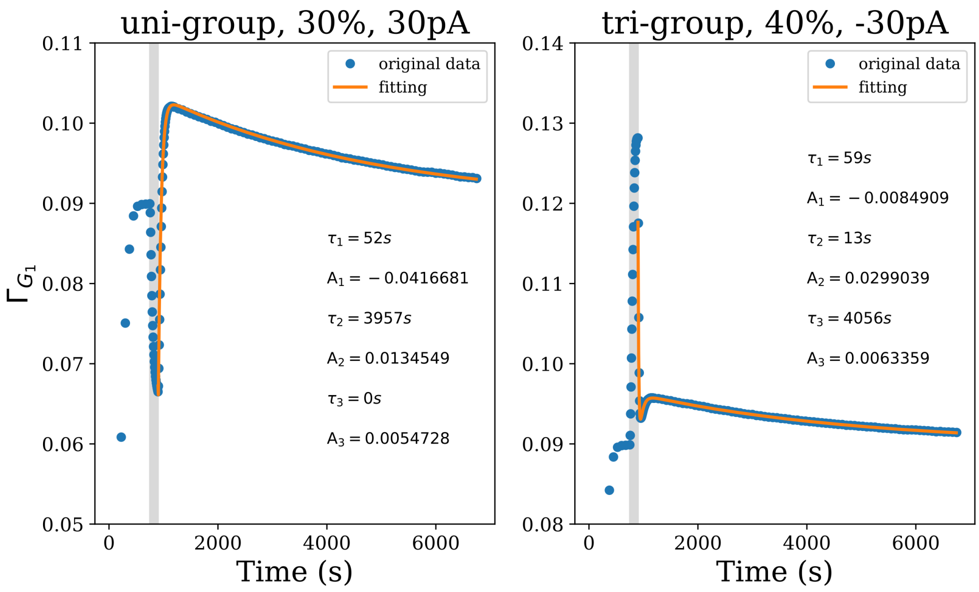


Figure S2: Examples of fitting a triple-exponential function. Left: Fitting the connectivity in the uni-group scenario, where $\tau_3$ amounts to 0 s. Right: Fitting the connectivity in the tri-group scenario. In both cases, the grey bar represents the stimulation phase; the time before the grey bar is the growth phase and after it comes the relaxation phase.
